# Sri Lankan Footprint in Global Aquatic Science Research: A Scientometric Analysis

**DOI:** 10.1101/2022.01.11.475849

**Authors:** Tharindu Bandara

## Abstract

The present study summarizes the research productivity and international collaboration in aquatic studies conducted by Sri Lankan scholars during 2000-2019. The study was based on the SCOPUS^®^ database. R programming language, package *bibliometrix* and Vosviewer software were employed in the analysis. Results of the present study indicate that increasing growth trend in the annual number of publications. A significant correlation (p<0.05) between the number of articles and per capita GDP (Gross Domestic Production) was also observed. Senior authors dominated in terms of the article count, citation count, *h* index, and other author productivity indices. Journal of the National Science Foundation of Sri Lanka had the highest article count (n=33). Aquatic studies in Sri Lanka were more locally funded. Sri Lanka had strong research collaborations with Japan, South Korea and Australia. During 2000-2019, the transition of aquatic studies from lacustrine field studies to molecular lab-based studies were observed. The findings of the present study may provide a comprehensive understanding on the current context and future directions of aquatic studies in Sri Lanka.

## Introduction

Sri Lanka, an island nation located in the Indian ocean comprised of more than 10,000 reservoirs and reported as the country with the highest reservoir density in the world (FAO, 1997). The reservoirs represent 75% of the inland water surface of Sri Lanka (Kularatne, Pascoe, Wilson, & Robinson, 2019) and the remnant is represented by riverine and coastal ecosystems. Over the years, these aquatic ecosystems provide valuable ecosystem services for the community and the importance of these systems were also highlighted in terms of national development. More recently, the sustainable utilization of these aquatic resources is also highlighted with the ample number of scientific studies (Edirisinghe, 2003).

Scientific studies are an imperative tool for elucidating the importance and mechanisms of natural ecosystems. The early scientific studies on Sri Lankan aquatic systems have driven by foreign scholars (Apstein, 1907) and later on, several Sri Lankan scholars have contributed to the aquatic science research. Although initial aquatic studies in Sri Lanka have focused on basic limnological studies (Apstein, 1907), Sri Lankan aquatic studies have diversified into various directions in later decades. These include various studies in aquaculture, oceanography, aquatic ecology and taxonomical studies. On the other hand, the national policies of Sri Lanka have given considerable attention to both freshwater and marine resources as one of the potential resources for elevating the country’s economy. On par with that, education on aquatic resources has been widened through dedicated undergraduate programs for aquatic sciences (Bandara & Radampola, 2017). Apart from that many universities offers a wide range of postgraduate course units related to the aquatic sciences. This, in turn, has generated a number of scholars who are contributing to national and international studies in the field of aquatic sciences.

Although there are several studies on aquatic resources in Sri Lanka, many of those studies are limited to national open access journals which are circulating among the local scientific community. Despite the development of the worldwide web and the indexing services, many of these Sri Lankan journals have not been indexed in popular databases such as SCOPUS^®^ and Web of Science™ (ISI) (science citation or social science citation indices). Currently, the only journal published in the science citation index-expanded is the Journal of the National Science Foundation of Sri Lanka. However, this does not imply that Sri Lankan research is outperformed in the international arena. Still today, a considerable number of publications are appearing in the above databases through reputed international journals. Nevertheless, there were no quantified studies that focus on the productivity of the aquatic studies with Sri Lankan institutional footprint. Quantified scientific output is essential for elucidating the contribution of aquatic resources for national development, understanding the conservation needs and identifying the research gaps. Although bibliometric studies are not direct measure for evaluating the science (Ivanović & Ho, 2014), it has been utilized for identifying trends in the particular scientific field (Bandara, 2020), evaluating and comparison of the authors (Bond, Clout, Czernkowski, & Wright, 2020; Singh, Mittal, & Ahmad, 2007). Therefore, the present work focused on i). To assess the Sri Lankan authors’ contribution to aquatic science research in SCOPUS^®^ database using bibliometric measures ii). Identifying the network structure of collaboration among authors iii). Elucidate the key themes in Sri Lankan aquatic studies and its’ evolution.

## Materials and Methods

### Data collection

Articles for the present study were collected from the SCOPUS^®^ database (www.SCOPUS.com) published by ELSEVIER (www.elsevier.com). The following search query was employed to retrieve the articles related to the aquatic studies by Sri Lankan institutional affiliations, during 2000-2019. Only articles written in the English language were utilized in the study. The asterisk wildcards used for the truncation and retrieving the various terms related to the main keyword (e.g., fish* (fisheries, fishes), Aqua* (aquatic, aquaculture)). The search was carried out on 15-June-2020.

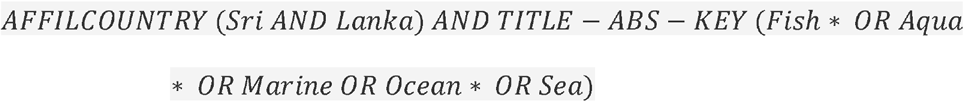

The above search query retrieved 874 documents. Articles were manually evaluated for title, abstract and keywords to find the mismatches. Articles related to groundwater and geology were omitted as those were not directly linked with the scope of the current study. Therefore, a total of 510 articles were utilized in further analyses.

### Data analysis

Metadata of the selected articles were exported in *.RIS* format. Bibliometric analyses were performed by using *bibliometrix* package (Aria & Cuccurullo, 2017), R programing language (R Development Core Team, 2017) and R studio integrated development environment (IDE) (RStudio Team, 2020). To determine the bibliographic network matrices, Vosviewer (version 1.6.11) visualization tool was employed (Van Eck & Waltman, 2010).

## Results and Discussion

### Growth trend of articles during 2000-2019

From 2000-2019, the number of articles related to aquatic studies has shown an increasing trend (R^2^ =0.72) (Figure 1). Since 2016, more than 40 annual publications appeared in the SCOPUS^®^ database. The lowest number of articles (n=5) appeared in 2004 and the highest number of articles appeared in 2017 (n=70).

**Figure 1:**
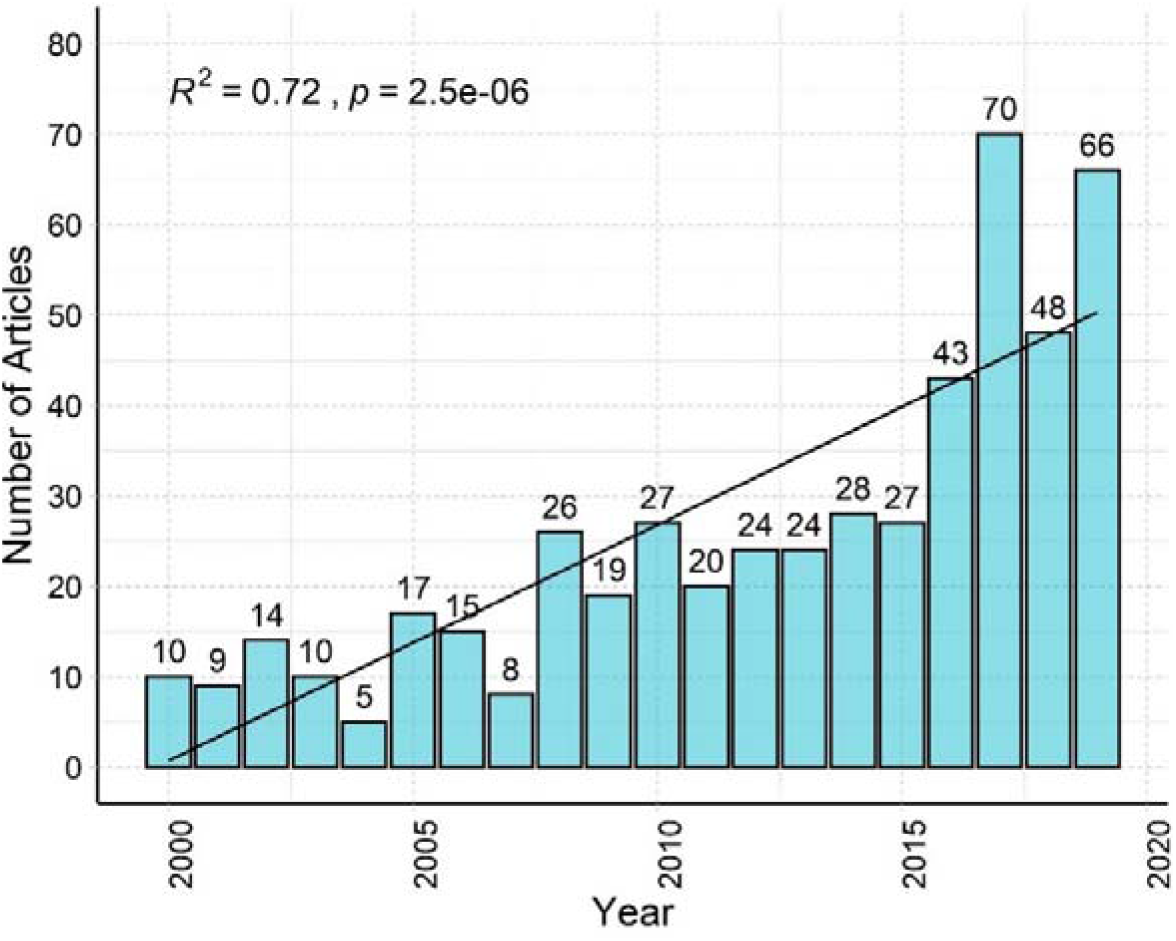
Number of articles appeared in the SCOPUS® database related to the Sri Lankan aquatic studies (2000-2019)

### Top 10 Journals

The top 10 journals with the highest number of publications were presented in Table 1. Journal of the National Science Foundation of Sri Lanka (JNSFSL) has published 33 articles related to aquatic studies. Among the top 10 journals, the lowest number of articles (n=6) have appeared in the Aquaculture Research, Ocean and Coastal Management, Marine Pollution Bulletin and the Asian Fisheries Science journals. To assess the impact of each journal Scimago Journal Ranking (SJR) was utilized. SJR was developed in 2007 (Bollen, Rodriquez, & Van de Sompel, 2006) based on the SCOPUS^®^ database and it measures journal’s prestige/influence based on the average number of weighted citations received in the selected year by the documents published in the same journal for last three years (González-Pereira, Guerrero-Bote, & Moya-Anegón, 2010). Higher the SJR Ranking, the journal has a higher influence/prestige in the given subject area. Although the JNSFSL published the greatest number of articles, the related SJR ranking was the lowest (0.139).

**Table 1:**
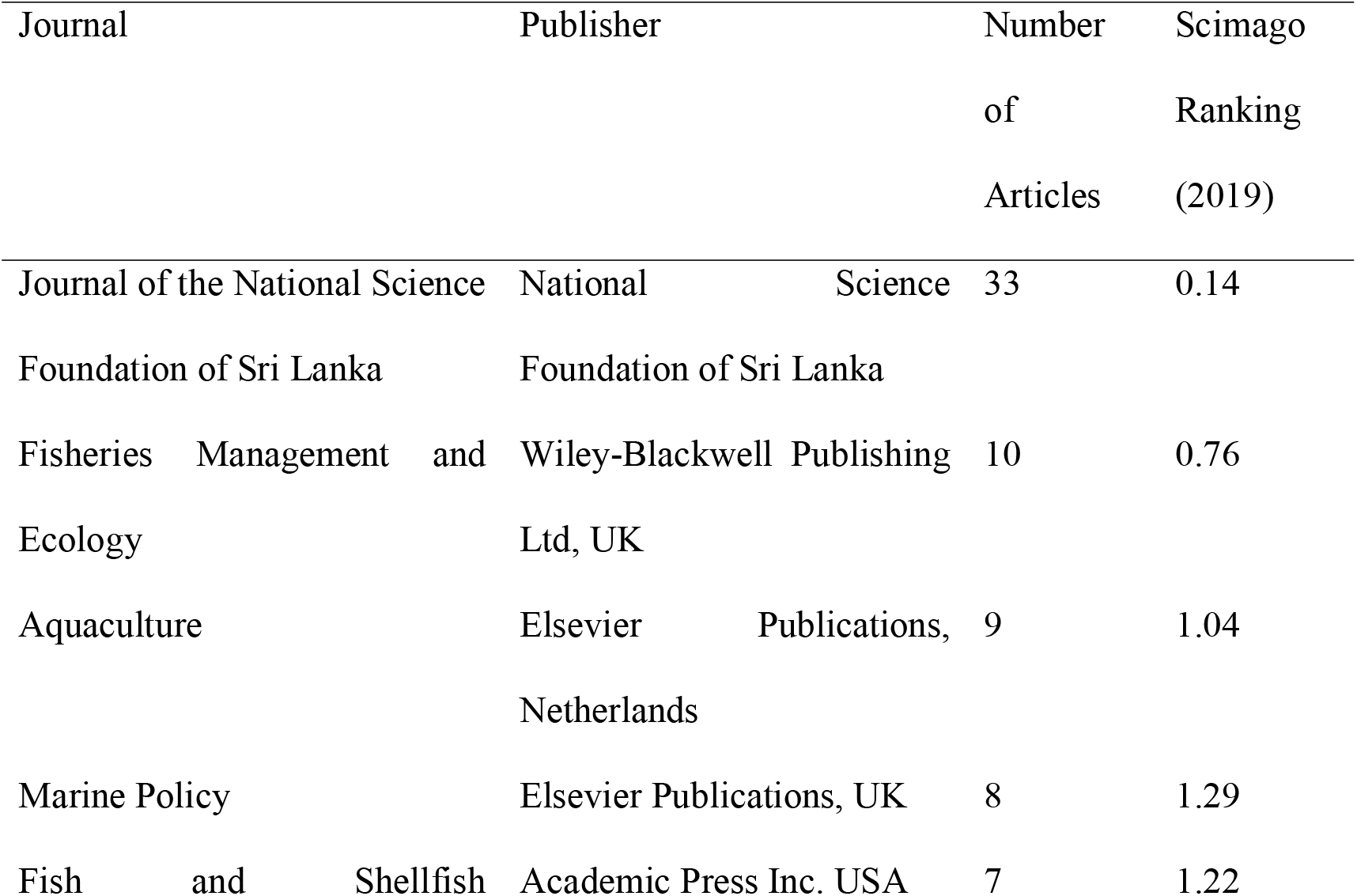
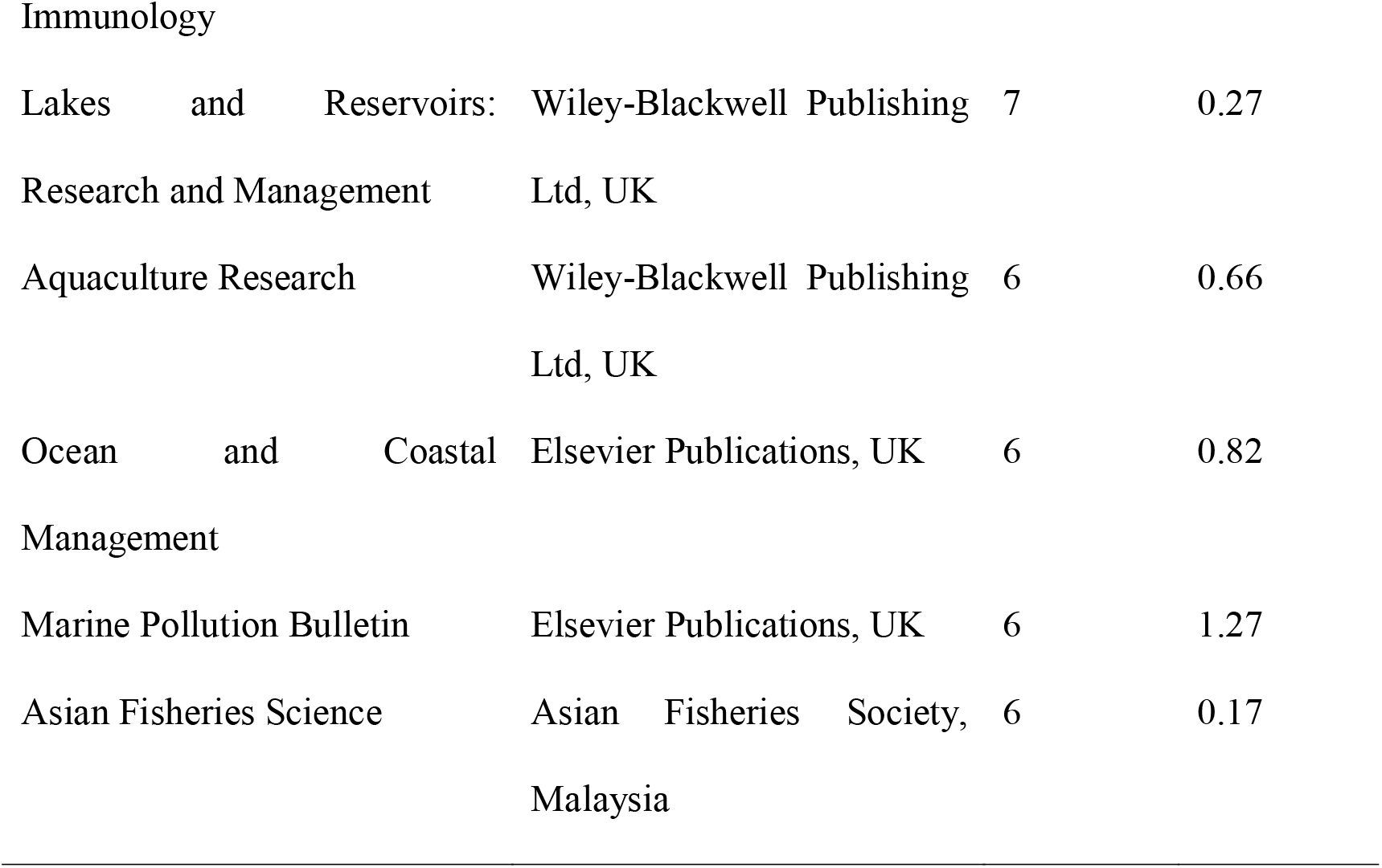
Top 10 journals with the highest number of articles, journal publisher and SJR

This might be attributed to the weighted score of SJR is more biased to the more prestigious international journals (Delgado-LÓPez-CÓZar & Cabezas-Clavijo, 2013; Falagas, Kouranos, Arencibia-Jorge, & Karageorgopoulos, 2008) than the national/regional journals. This low SJR is also apparent in the regional journal; Asian Fisheries Science. JNSFSL is the first Sri Lankan Journal indexed in both SJR and Web of Science™ database. Moreover, it has no article processing fee/publication fee. The higher number of articles appearing in this journal might be advocated the authors’ interest in publishing articles that has more national importance and concerns over the publication cost in gold or hybrid open access journals. Therefore, JNSFSL might be a better option for more authors to publish their research. Conversely, Marine Policy has reported the highest SJR ranking (1.29) among the selected journals and it covers the broader area of coastal and marine policies, management practices, national, regional, international marine etc.

### Authorship pattern

During 2000-2019, a total of 1523 authors participated in aquatic science research. However, the actual number of authors with Sri Lanka affiliation is lower than that, since most of the research papers are multi-authored. Among 510 articles, 32% of articles represent Sri Lankan corresponding authorship. Figure 2 illustrates the top 10 authors productivity over time. Amarasinghe US has the highest number of articles (n=37). Amarasinghe US and DE Silva SS have published their articles continuously from 2000-2019. Jayathissa LP is the most cited author (n_total_= 373) is followed by Amarasighe US (n_total_= 329) and DE Silva (n_total_= 321).

**Figure 2:**
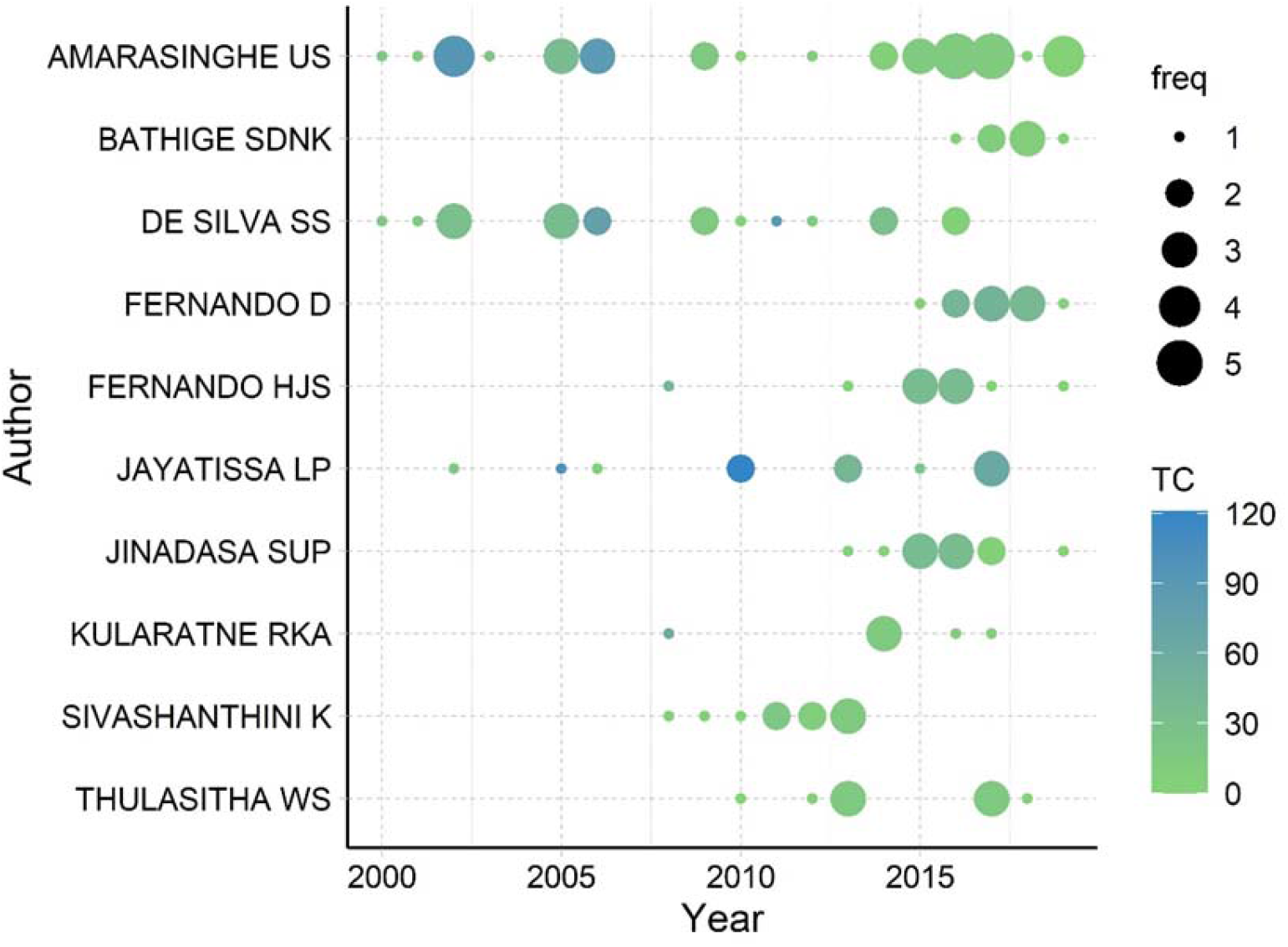
Top 10 authors (based on the total number of articles) productivity over time (TC=Total number of citations for the respective year, freq=number of articles published in the respective year)

### Impact of the author

Based on the SCOPUS^®^ data, the author impact measured by various indices has presented in Table 2. Hirsch index (*h*-index) usually utilized to quantify the authors’ scientific output (Andres, 2009). By definition, “A scientist has index *h* if h of his or her Np papers have at least h citations each and the other (Np - h) papers have fewer than h citations each” (Hirsch, 2005). Over the years *h*-index considered as the aggregated research contribution of a researcher and utilized to indicate eminence and comprehensive impression of an author (Khosrow-Pour, 2017). On the other hand, the *g*-index is an improvement of the *h*-index as proposed by Egghe (2006). It avoids the disadvantage of the *h*-index which underrate the most cited papers. As suggested by Egghe (2006), *g*-index accounts for the most cited papers in the calculation and as a rule of thumb, g ≥ h. Therefore, authors with a higher *g*-index may have more highly cited papers with more international impact (Andres, 2009). Amarasinghe US was the top author with the highest *h* and g-indices. The *m*-index/m-quotient is the ratio between the *h*-index and the number of years elapsed since the author’s first publication (Khosrow-Pour, 2017). Therefore, authors who have started to publish recently may have a higher *m*-index. The highest *m*-index is reported by Fernando D whose first publications appeared in 2015. In contrast, authors with a longer career since the first publication may have a lower *m*-index. This was evident from the *m*-indices of the more senior authors; Amarasighe US; De Silva SS and Jayathissa LP etc. Conversely recorded low value of the *m*-index by some of the authors (e.g. Thulasitha WS) is primarily due to a lower *h* value than the elapsed time since the first publication. However, the indices presented in Table 2 might be different from the actual indices since the data in the present study focused only on the SCOPUS^®^ database. Moreover, the study did not count the other sources (e.g. book chapters, conference proceedings etc.) for the calculation of the above indices.

**Table 2:** Impact of the top 10 authors based on major productivity indices

| Author | $h$-index | $g$-index | $m$-index | Year of first publication in |
| --- | --- | --- | --- | --- |
|  |  |  |  | SCOPUS<sup>®</sup> database |
| AMARASINGHE US | 10 | 17 | 0.476 | 2000 |
| DE SILVA SS | 9 | 17 | 0.429 | 2000 |
| PATHIRATNE A | 9 | 15 | 0.429 | 2000 |
| JAYATISSA LP | 8 | 10 | 0.421 | 2002 |
| FERNANDO D | 6 | 10 | 1 | 2015 |
| FERNANDO HJS | 6 | 10 | 0.462 | 2008 |
| JINADASA SUP | 5 | 8 | 0.625 | 2013 |
| SIVASHANTHINI K | 5 | 6 | 0.385 | 2008 |
| THULASITHA WS | 5 | 5 | 0.455 | 2010 |
| BATHIGE SDNK | 4 | 4 | 0.8 | 2016 |

### Scientific productivity

The scientific productivity of a given field can empirically be evaluated using Lotka’s law. In general Lotka law specifies the number of authors contributing *n* contributions is approximately about those making 1/n^a^, where a=2 (Coile, 1977). Lotka’s law of author productivity was applied to figure out the scientific productivity of the articles in aquatic studies with Sri Lankan institutional affiliation. Scientific output related to the aquatic studies research with Sri Lankan affiliation has indicated that the β coefficient of 2.39 and a constant of 0.32 with Kolmogorov-Smirnoff goodness-of-fit test of 0.88 (p=0.04, two-sample t-test) (Figure 3). This discrepancy between the theoretical and observed Lotka distributions has indicated that aquatic studies affiliated with Sri Lanka have followed the Lotka law.

**Figure 3:**
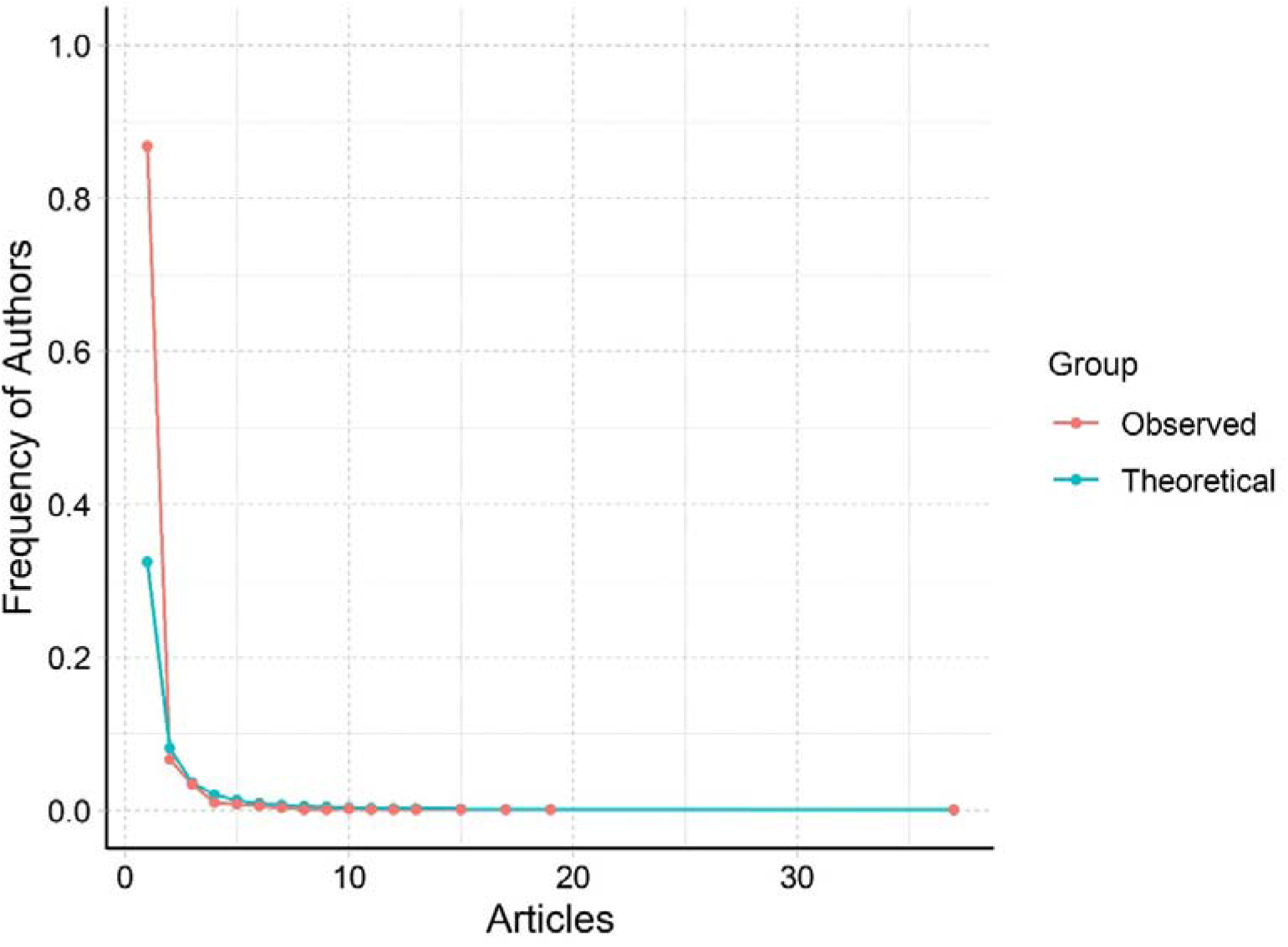
Author productivity based on Lotka’s law.

### Publication output and per capita GDP

Research and development of a country is strongly ties with its economic conditions and usually, high-income countries are frontiers in high-quality publications. This may evident in many indices such as a higher number of citations per paper, higher per capita publications (Hazelkorn, Coates, & McCormick, 2018) in developed countries. Gross Domestic Production (GDP) is an economic measure used to evaluate the economic strength of a country measured in terms of total services and goods of a country in a given year. Rather than the GDP, improvements of the per capita GDP ensures economic well being and technological improvements of a country (Peterson, 2017). Several studies have been conducted to understand the relationship between the GDP and publication output of a country as well as the per capita GDP and publication output of a country (Cheng & Zhang, 2013; Liang et al., 2015; Luo et al., 2015). In the present study, a significant positive relationship between per capita GDP and the number of articles was revealed (Figure 4). In contrast, to present findings, Meo, Al Masri, Usmani, Memon, and Zaidi (2013) found no significant relationship between per capita GDP and the number of articles in a study to find out the impact of GDP on the scientific research in Asian Countries. This may due to discrepancies between the GDP and the Gross Expenditure in Research and Development (GERD) in many Asian countries. Although it is advisable to use more direct indices such as GERD for the understanding of the research productivity, data on GERD of Sri Lanka were sporadically available. However, for more conclusive results it would be advisable to gather the data from other databases (e.g. Web of Science™).

**Figure 4:**
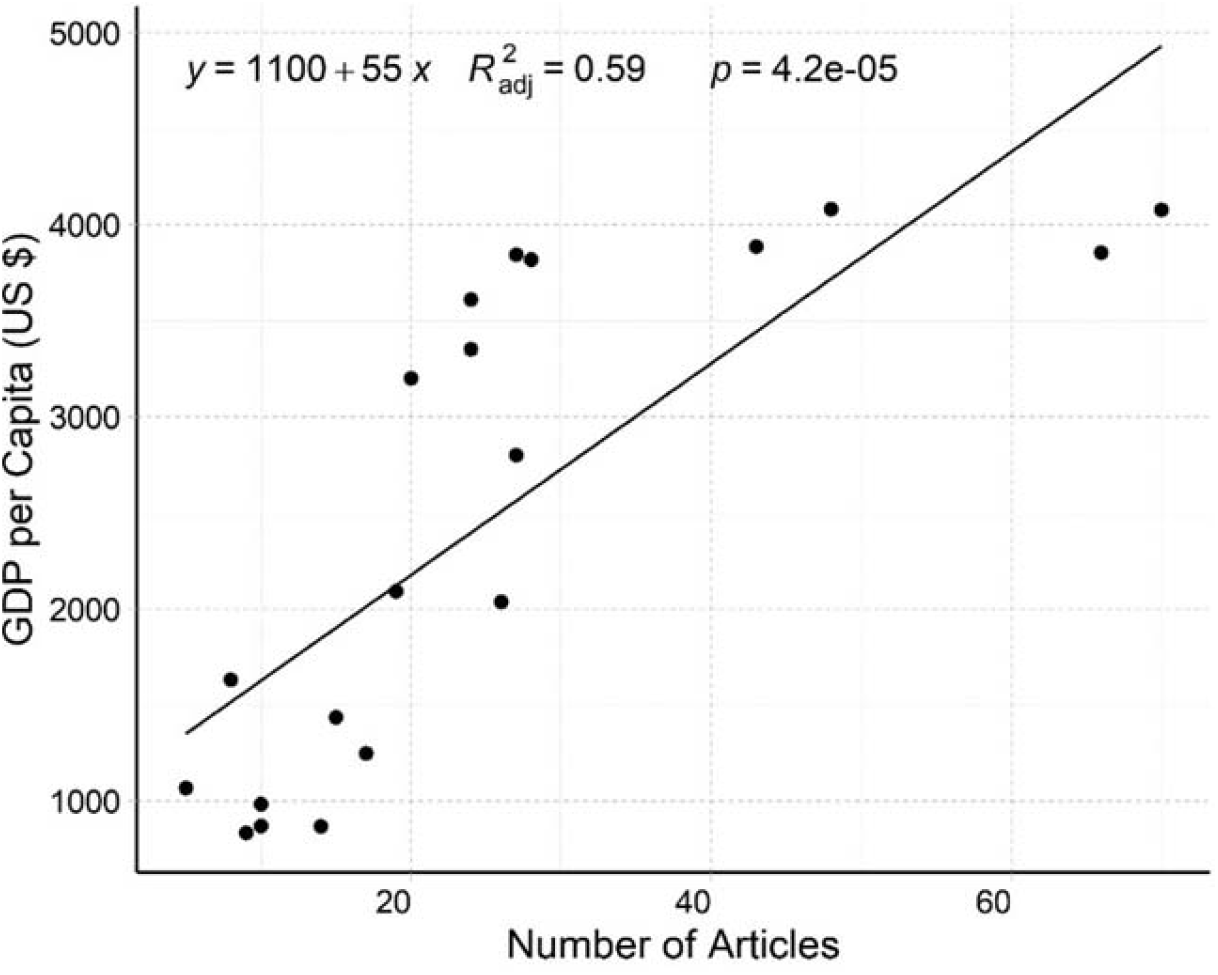
The relationship between the per capita GDP and the number of articles.

### Funding

Among the total number of publications, 104 articles have indicated the funding information. These fundings include early-career research grants (e.g. PhD), project grants, mobility grants etc. Analysis of the funding details indicated that internal fundings (Sri Lanka) have contributed to the most number of studies in aquatic research (n=29) (Figure 5). The major funding agency for internal research is the National Science Foundation of Sri Lanka.

**Figure 5:**
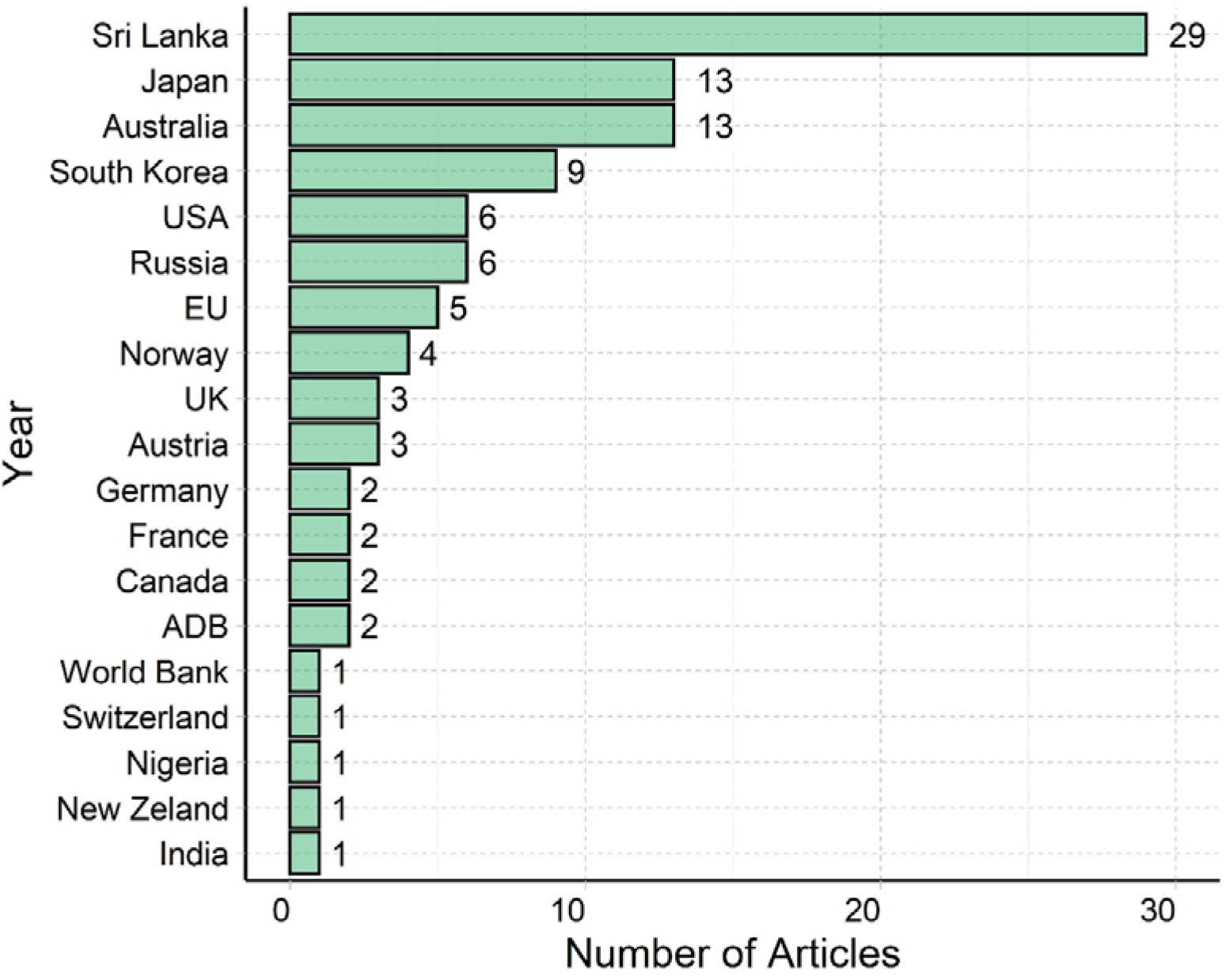
Number of publications funded by various countries (EU-European Union direct funds; ADB-Asian Development Bank)

However, The total number of international research funding (n=75) has surpassed the number of internal fundings. The unavailability of the monetary values of these fundings has impaired the country based total monetary value calculations. As a lower-middle-income country where budgetary austerity predominates, international funding may provide additional support for Sri Lankan aquatic studies research. In this scenario, international fundings are important in optimizing the aquatic research and align the research with national priorities (e.g. eliminating poverty). Moreover, these international funds are necessary for project implementation and training of early-career researchers. Previous research has indicated that international funding may have a higher publication impact (Wang & Shapira, 2015). Japan and Australia are important destinations for Sri Lankan scholars for their higher studies (e.g. PhD). Annually, several scholarships are offered to Sri Lankan scholars from these countries to pursue their higher studies (e.g. MEXT scholarship, Japanese Government Monbukagakusho Scholarship, Endeavour scholarship and Australia awards etc.). In recent years, South Korea was an emerging destination for Sri Lanka scholars to pursue their postgraduate studies and collaboration with Sri Lankan universities and South Korean universities are also emerging.

### Top 10 cited papers

Based on the total number of citations the top 10 articles has represented in Table 3. Analysis of these articles indicated that articles address multiple research areas including natural disasters (Tsunami), environmental toxicology, marine biology etc. Among the top 10 articles, the article entitled “chronic renal failure among farm families…” addressed one of the key issues in Sri Lankan agriculture sector. Elevated dietary Cadmium is highly debated for chronic renal failure in the north-central province of Sri Lanka and many studies are still focusing on this issue (Ananda Jayalal, Jayaruwan Bandara, Mahawithanage, Wansapala, & Galappaththi, 2019; Bandara, Wijewardena, Liyanege, Upul, & Bandara, 2010). Moreover, after the Tsunami catastrophe in 2004, several ecological research has emerged in evaluating the importance of the mangrove ecosystem on the island.

**Table 3:**
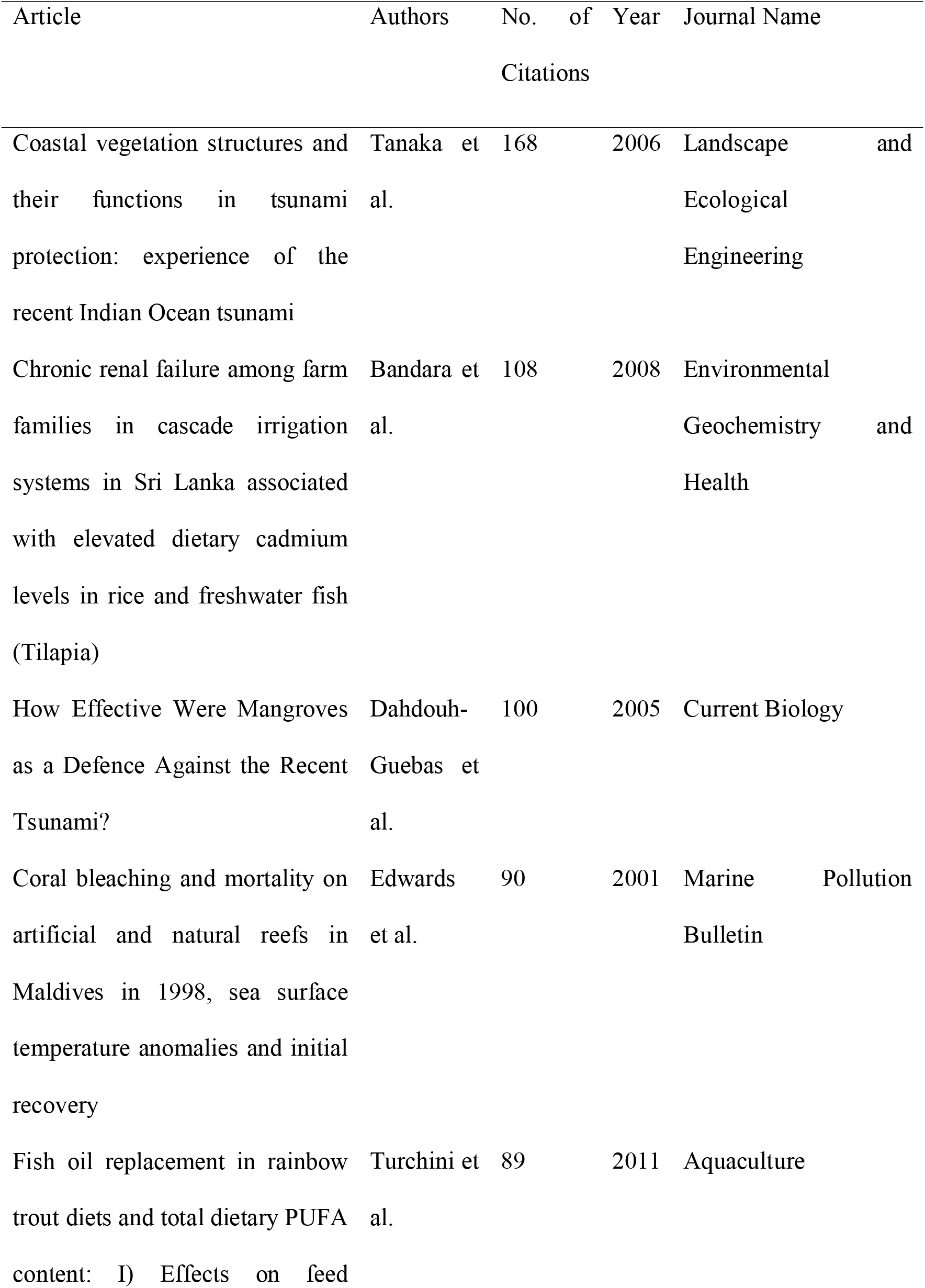
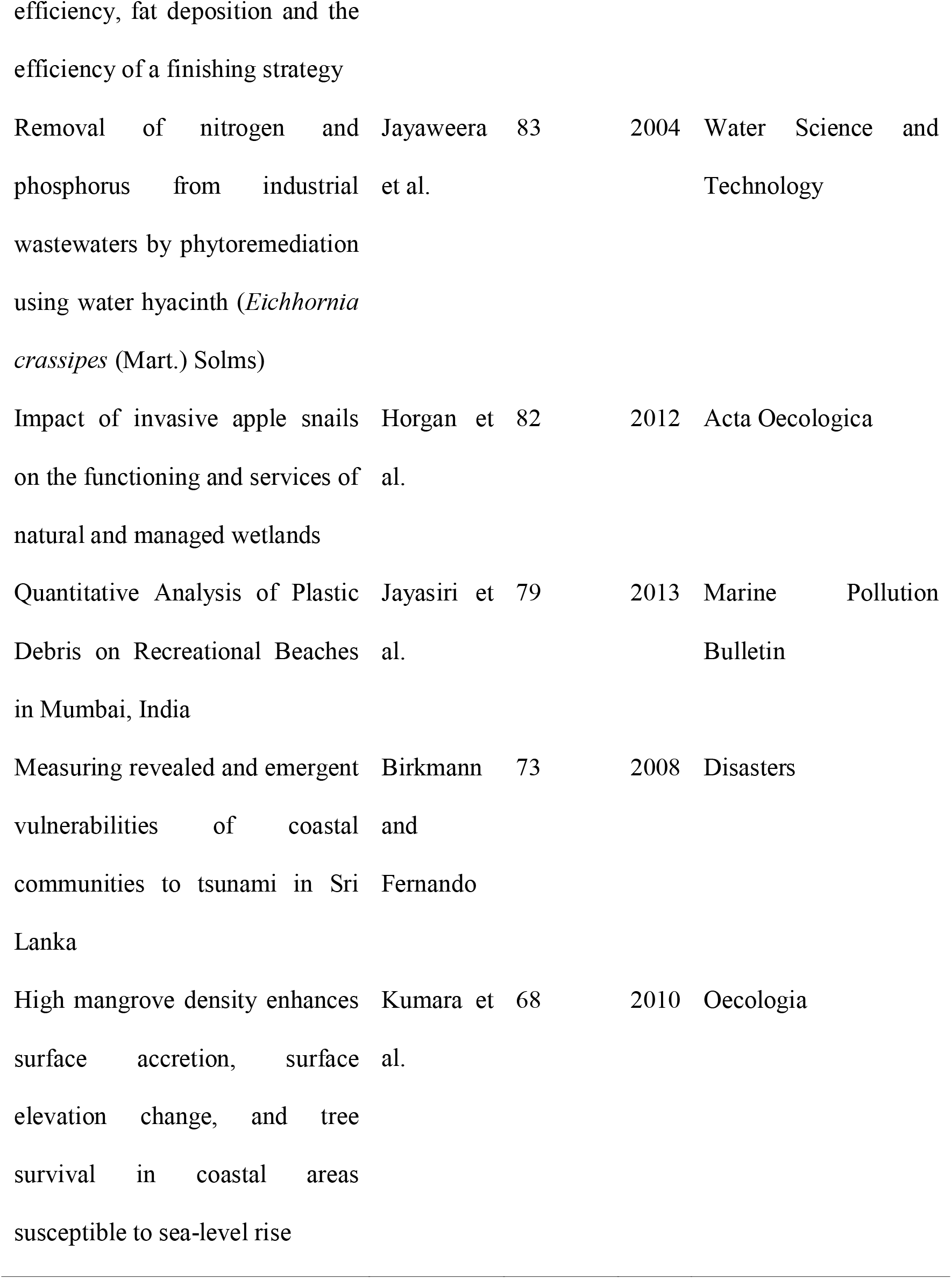
Top 10 highly cited papers

### Article distribution by institutes

The distribution of articles by institutes has illustrated in Figure 6. The University of Peradeniya has the highest number of articles (n=128) which is followed by the University of Ruhuna (n=79) and the University of Kelaniya (n=75). Several international institutes and universities were also evident in the results as the authors of these publications affiliated to multiple institutes. The University of Peradeniya is the largest university of Sri Lanka and hosts several postgraduate institutes and faculties facilitating aquatic studies research. Faculty of Fisheries and Marine Science and Technology, University of Ruhuna, Sri Lanka is a dedicated faculty for fisheries and marine science research in Sri Lanka. This may imply the highest number of articles from the above institutions. The author with the highest number of articles (Amarasinghe US) was affiliated with the University of Kelaniya and may have a significant impact in uplifting the article count in the same institution.

**Figure 6:**
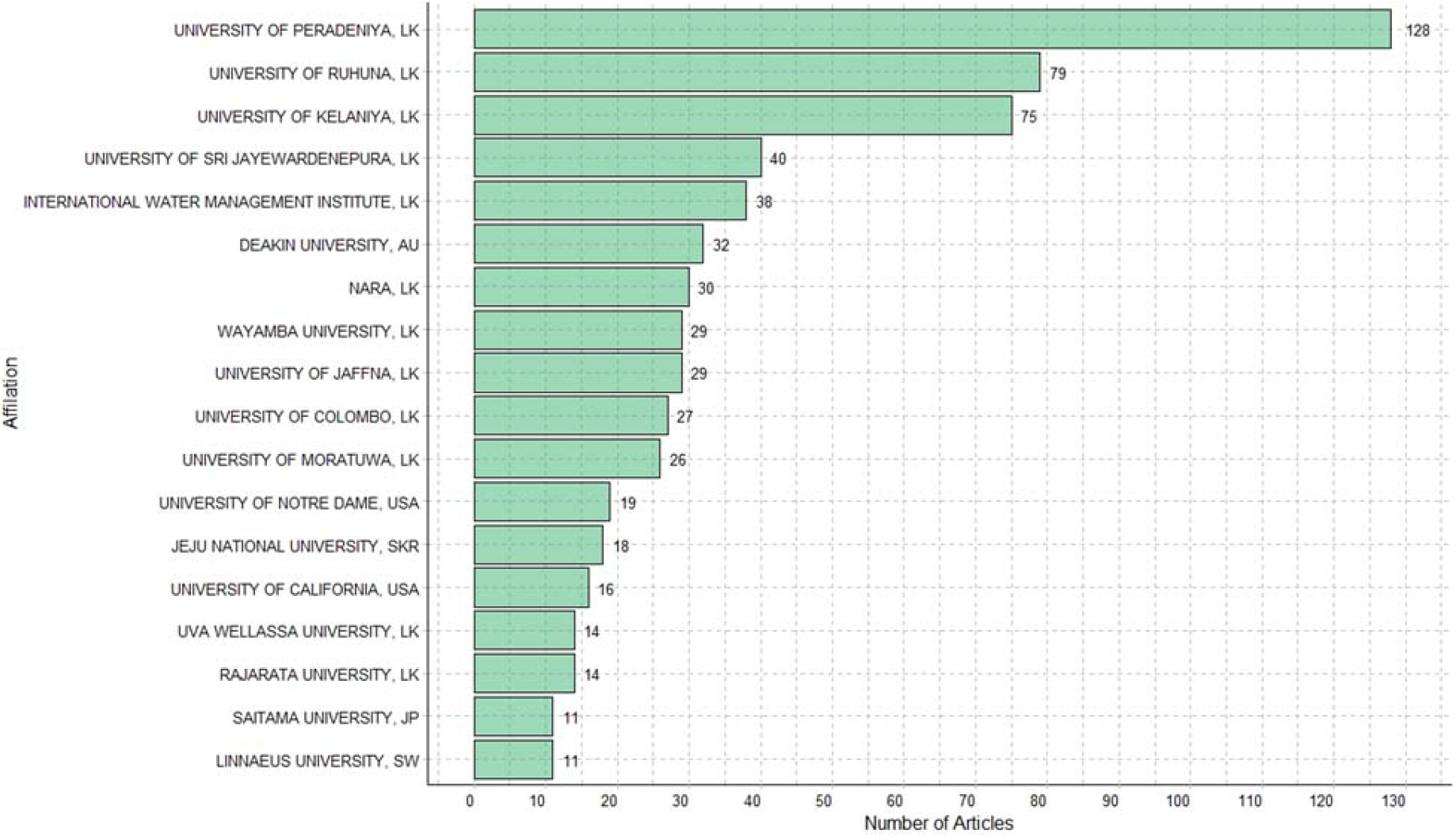
Article distribution by institutes. (LK: Sri Lanka, AU: Australia, SKR: South Korea, JP: Japan, SW: Sweden)

### Thematic map

Evaluation of the author keywords in the strategic diagram has illustrated in Figure 7. The strategic diagram is based on the density and centrality of major themes plotted along the x and y-axis (x axis-centrality; y axis-density) (Cobo, López-Herrera, Herrera-Viedma, & Herrera, 2011). The strategic diagram consists of four different quadrants (Cobo et al., 2011).

**Figure 7:**
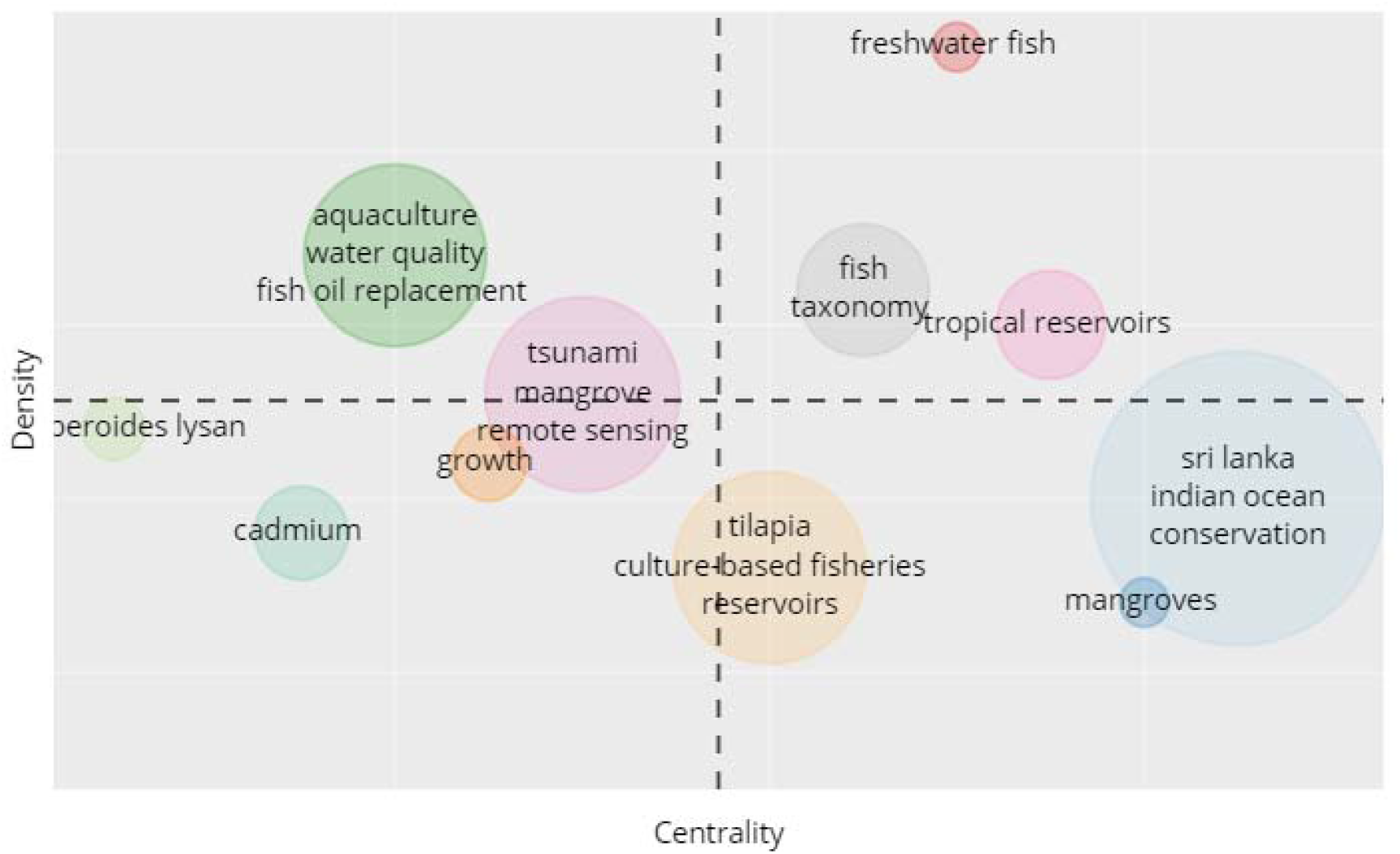
Strategic diagram based on the number of author keywords.

1. Themes in upper left quadrant: Highly developed and isolated/specialized themes
2. Themes in upper right quadrant: Motor themes (themes in this groups are conceptually related to the other themes)
3. Themes in lower left quadrant: Emerging or declining themes
4. Themes in lower right quadrant: Basic and transversal themes

Based on the above classification, themes related to aquaculture nutrition can be identified as highly specialized theme. This includes the narrow research area as denoted by fish oil replacement. Studies related to the tropical reservoirs and fish taxonomy were motor themes. As discussed earlier, studies related to Cadmium or chronic kidney disease in the north-central province of Sri Lanka is one of the emerging research areas. With technological advancements, studies related to the Geographic Information Systems (GIS) and remote sensing has gained much attention from Sri Lankan scholars. On the other hand, basic and transversal themes contained studies on mangroves and conservation of aquatic resources related themes.

### Network analysis

#### Co-authorship

Results of the co-authorship analysis have clustered the authors (*n*=16) into 5 distinct clusters (Figure 8). The size of the vertices as indicated by the loops represent the number of papers published. Cluster 1 consisted of the highest number of authors (*n*=5) is followed by cluster 02 (*n*=4). The size of each node is proportional to the number of papers published by each author and the width of the link between two nodes represent the intensity of co-authorship. Thicker the link between two nodes represents strong collaboration (Swar & Khan, 2014). Turchini GM dominates in cluster 1. Amarasinghe US holds the central position in cluster 2 and De Silva SS in cluster 3. The co-authorship intensity between Amarasinghe US and De Silva SS was the highest. Clustering of authors may also indicate the research areas of the authors. A detailed study on authors in cluster 1 indicates that aquaculture nutrition is the key research area of those authors. Cluster 2 represented the inland reservoirs and fisheries management while cluster 03 denotes aquaculture, inland fisheries and reservoir management. Authors in cluster 04 focus on culture-based fisheries, inland fisheries, biodiversity and related themes. Cluster 05 denotes environmental management and fisheries. However, the above demarcations may not be robust as the cliquishness of cluster 02-05 is higher than that of cluster 01.

**Figure 8:**
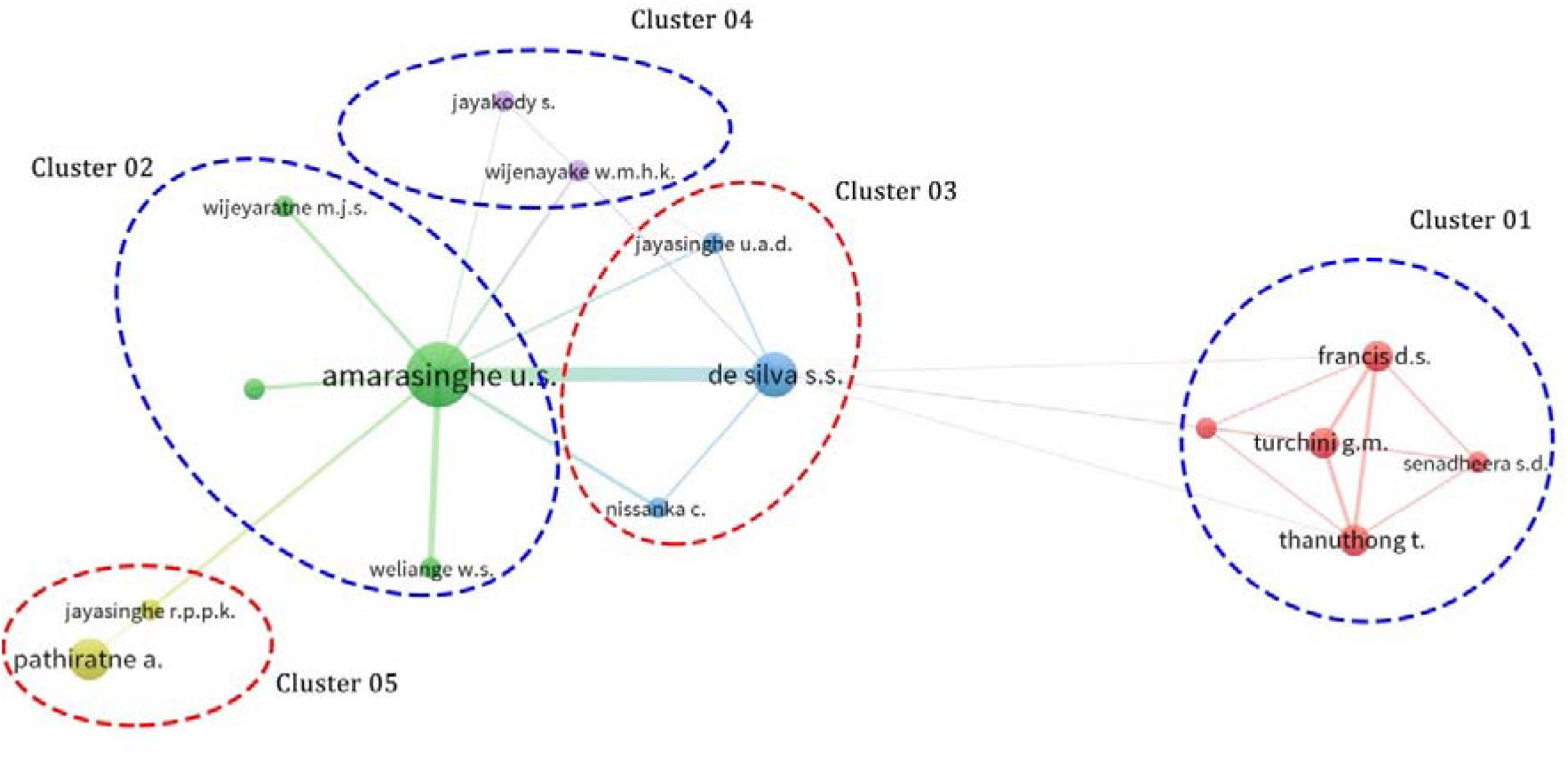
Co-authorship network of aquatic studies by authors with Sri Lankan affiliations (2000-2019)

#### Country-level co-authorship

Country-level co-authorship analysis indicates the research communication between the two countries. A country-level co-authorship network related to Sri Lankan aquatic studies has illustrated in Figure 9. The bigger the size of the node, the bigger the total number of articles or it indicates more influential countries. Increased width between two nodes more co-authored documents between the two countries. Sri Lanka represents the hub of the network with the highest number of articles. As indicated by the thickest node, the co-authorship between Sri Lanka and Japan is the highest among other countries. Sri Lanka-South Korea, Sri Lanka-Australia and Sri Lanka-Canada also had strong ties in terms of co-authorship. Overlay visualization of the country level co-authorship (Figure 10) indicated that Sri Lanka has started strong collaborations with China and South Korea recently. In contrast to that, during 2012-2015 Sri Lanka has more collaborations with Australia, Japan and Canada.

**Figure 9:**
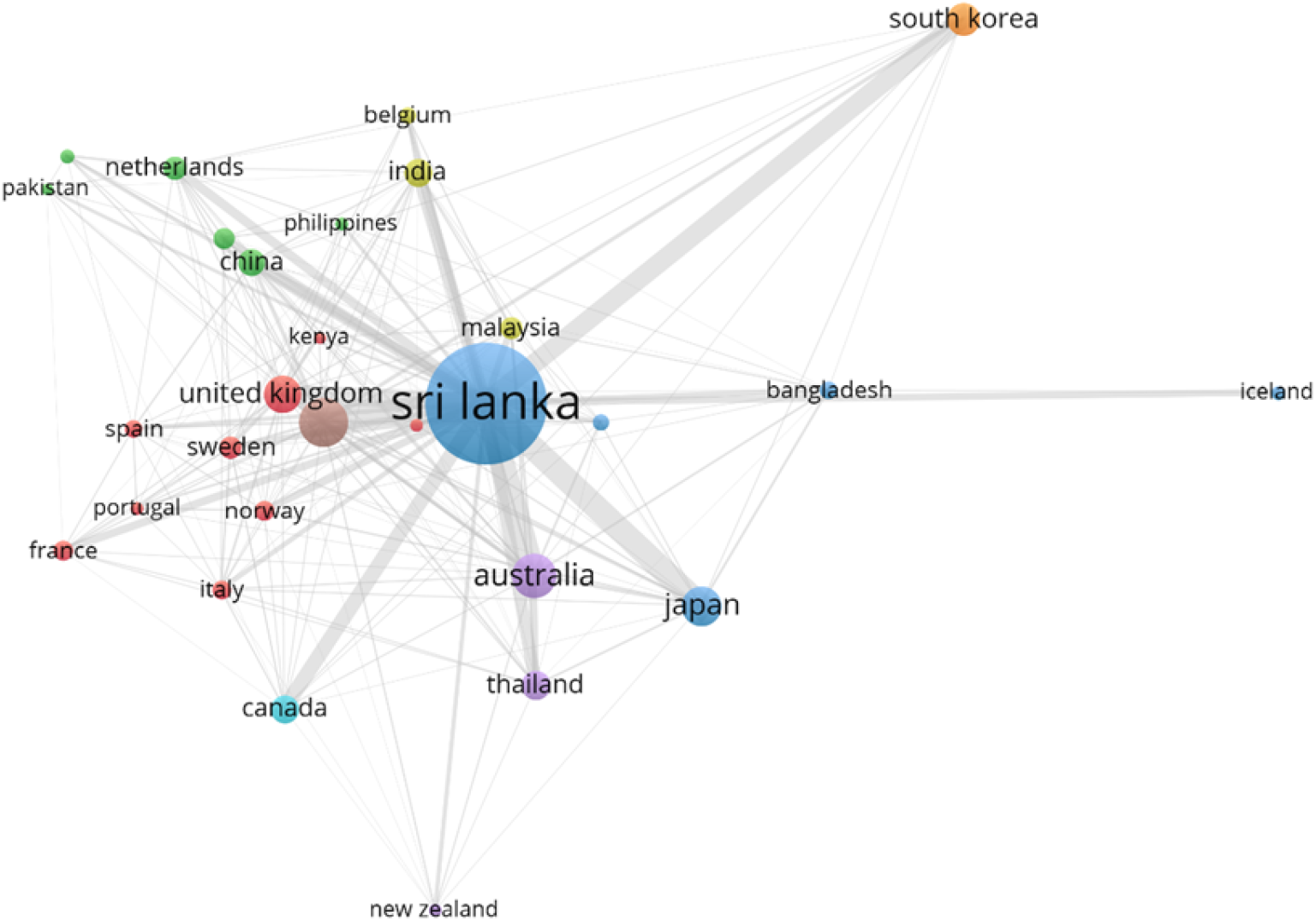
Country-level co-authorship network (2000-2019)

**Figure 10:**
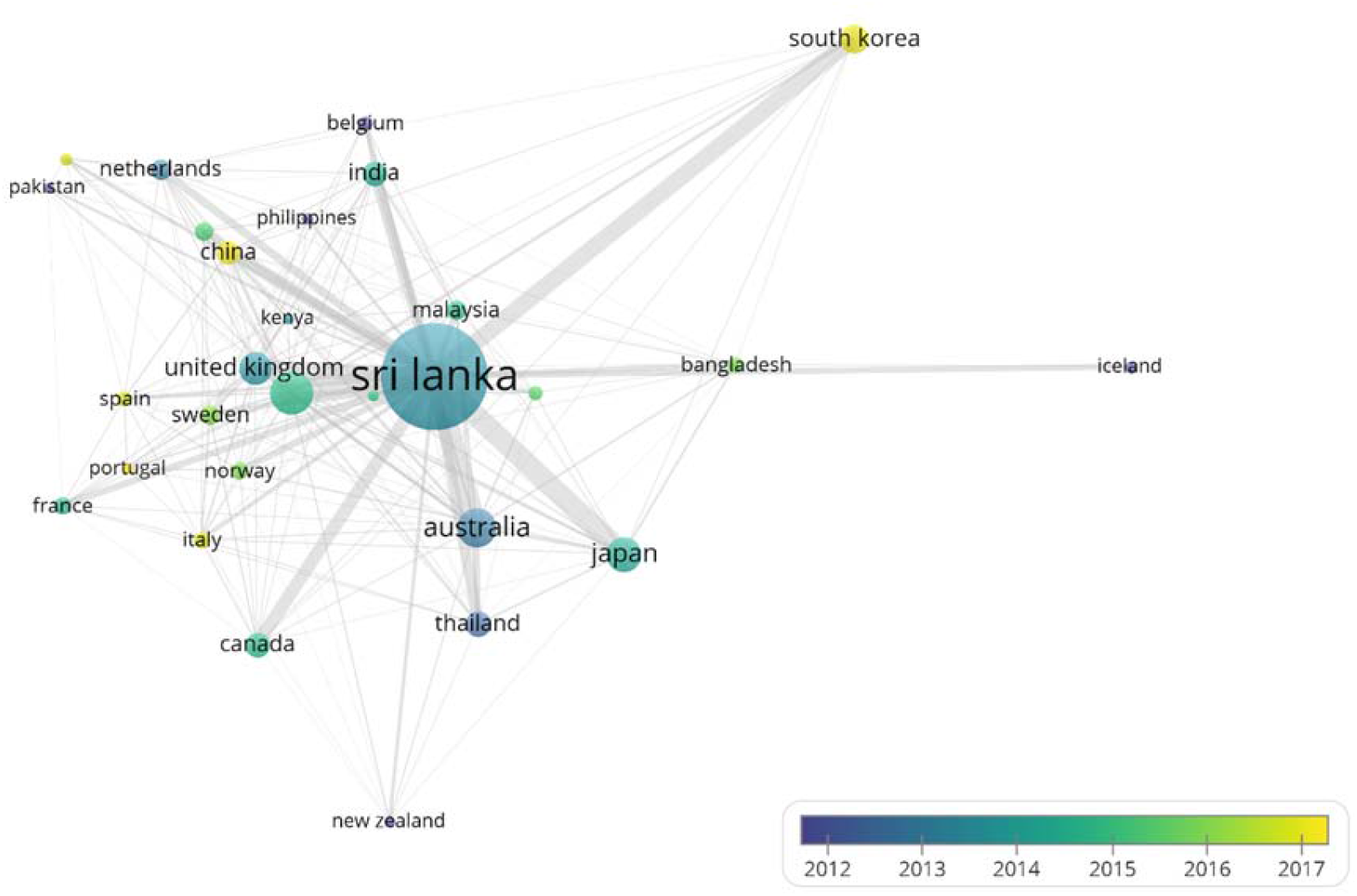
Overlay visualization of country-level co-authorship network (2000-2019)

#### Keyword co-occurrence

The keyword co-occurrence network has illustrated in Figure 11. The size of the node indicates the weights of the keywords. Bigger nodes have a higher weight (Liao et al., 2018). The thickness of the line between the two nodes is proportional to the co-occurrence of two words (Gu, Li, Li, & Liang, 2017) and link strength (*l_s_*) of a keyword indicate the frequency of co-occurrence (*f*) (Liao et al., 2018). Nodes that are close to each other have a higher relationship. The keyword co-occurrence diagram has depicted 5 clusters with different colours. Words in the same cluster are more related to each other. Cluster 01 consist of 27 keywords centred around the term *non-human* (*f* = 67, total *l_s_ =* 67), Cluster 2 was centered around *aquaculture* (f = 28, total *l_s_* =21). Cluster 3 was represented by *Sri Lanka* (f = 212, total *l_s_* =186). Cluster 4 was centred around the term *fish* (f = 56, *l_s_* = 57) and cluster 05 was represented by *water pollutants, chemical* (f = 28, *l_s_* = 28). The terms of cluster 01 represent the terminology related to molecular biology. Cluster 02 represent by the aquaculture studies in Sri Lanka. In cluster 03, the word Sri Lanka is strongly co-occurred with the fishery management, Tsunami and the reservoir (as represented by thicker lines). This has indicated that most of the aquatic studies in Sri Lanka have followed the research direction in fisheries management and reservoir fisheries. Co-occurrence of word fish with other related keywords in cluster 03 (*Oreochromis niloticus*, algae, growth rate) indicate research direction on the growth studies of tilapia. In cluster 05, water pollutant is more co-related with environmental monitoring, analysis, pollutant and exposure. This cluster has represented the research direction in environmental studies.

**Figure 11:**
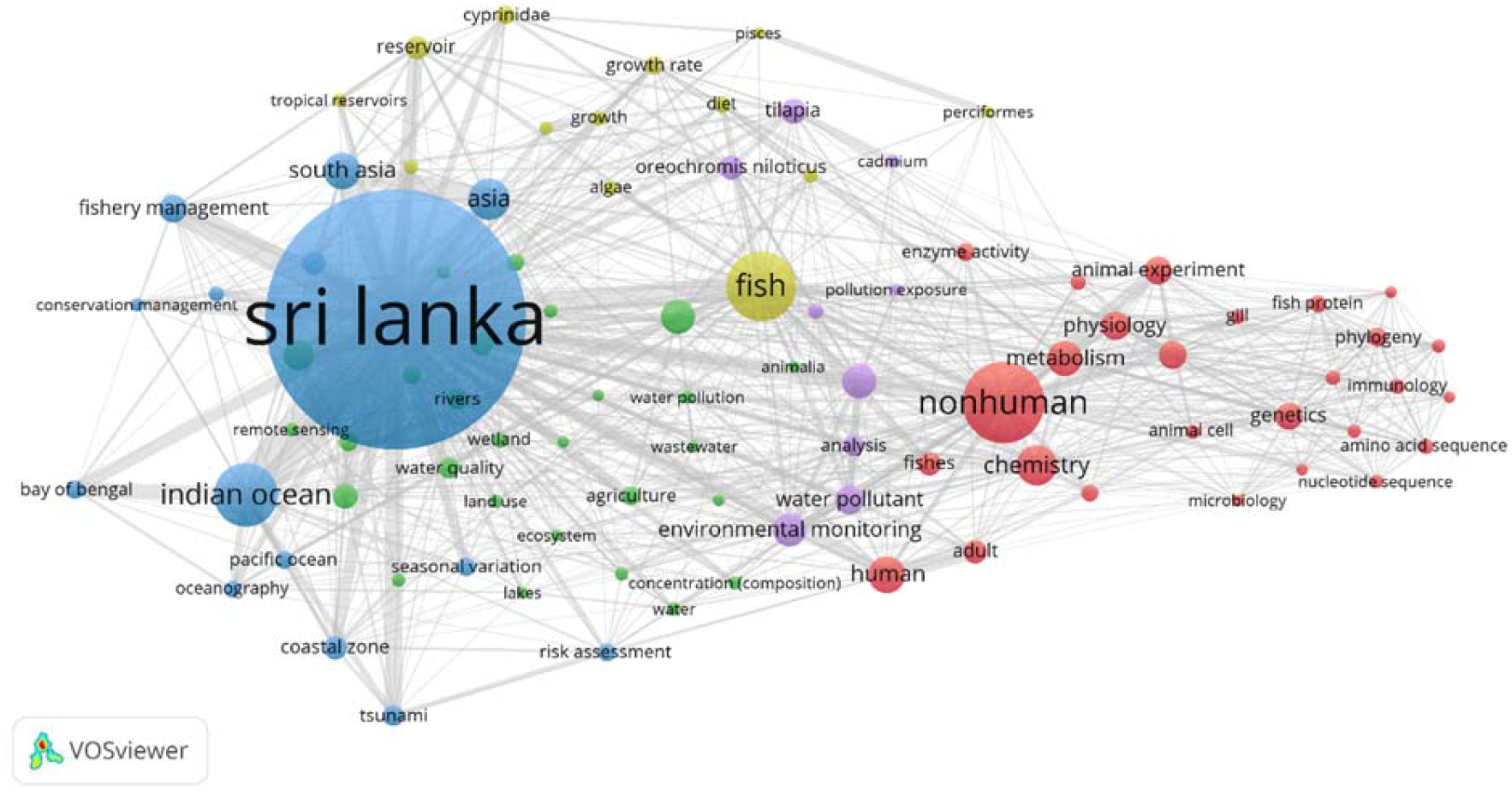
Keyword co-occurrence network of aquatic studies in Sri Lanka (2000-2019)

Overlay visualization of the keywords indicates the evolution of the aquatic studies research (Figure 12). During 2010-2012, studies were more focused on growth studies of tilapias and fisheries management. Since 2016 Sri Lankan authors affiliated studies were more focused on the molecular studies and studies in the Indian ocean, conservation and management of aquatic resources.

**Figure 12:**
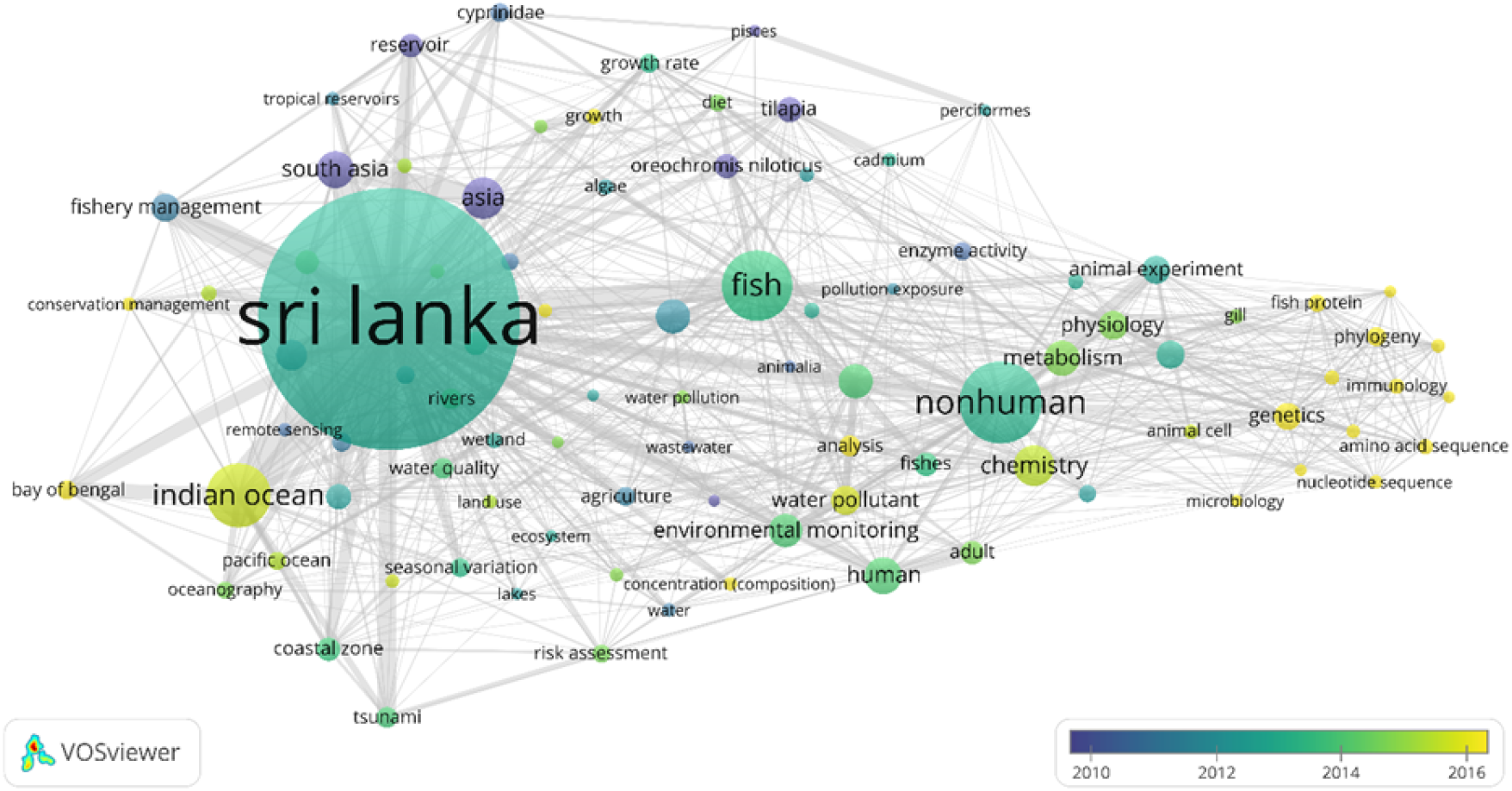
Overlay visualization of keywords related to the aquatic studies in Sri Lanka (2000-2019)

## Conclusion

The present study highlighted the key findings in Sri Lankan aquatic studies in a global context. The annual number of publications related to the aquatic studies are increasing in each year and it is strongly correlated with the per capita GDP. Despite many international journals, the Journal of the National Science Foundation of Sri Lanka (JNSFSL) holds a pivotal role in the number of articles published. This higher number of article count in JNSFSL may indicate more locally oriented research published by the Sri Lankan authors. More senior authors have a strong impact on aquatic studies in Sri Lanka and their affiliated institutions have strong rankings in terms of a number of publications. This may evident in the associated author productivity indices such as *h*-index. Authorship pattern of the articles has indicated that a large number of articles are multi-authored and authors with Sri Lankan affiliations also held the other affiliations concurrently. Asia-Pacific countries (Japan, South Korea and Australia) dominated in financial support for many studies affiliated by Sri Lanka authors. Evolution of the key research themes indicates that since 2016, more studies have focused on molecular studies than the fish growth studies and environmental studies. Network analysis of bibliometric data indicated that the strong collaboration of more eminent authors.

There were certain limitations in the present study. The number of publications may exceed the current amount as the study not focused on the other scholarly databases such as Web of Science™. Thus, it avoids the other related literature (e.g. books, proceedings etc.) and only concerened the articles published in the English language. Moreover, funding information of the articles were obscure in a number of articles and this may hinder the detailed analysis of funding related information. Therefore, future opportunities exist in extending this study considering these limitations for a more comprehensive outlook.

## Acknowledgements

The author wishes to acknowledge two anonymous reviewers for their constructive outlook on the manuscript.

## Conflict of Intrest

The author declare no conflict of intrest.

